# Information Processing by Simple Molecular Motifs and Susceptibility to Noise

**DOI:** 10.1101/023697

**Authors:** Siobhan McMahon, Oleg Lenive, Sarah Filippi, Michael P.H. Stumpf

## Abstract

Biological organisms rely on their ability to sense and respond appropriately to their environment. The molecular mechanisms that facilitate these essential processes are however subject to a range of random effects and stochastic processes, which jointly affect the reliability of information transmission between receptors and e.g. the physiological downstream response. Information is mathematically defined in terms of the entropy; and the extent of information flowing across an information channel or signalling system is typically measured by the “mutual information”, or the reduction in the uncertainty about the output once the input signal is known. Here we quantify how extrinsic and intrinsic noise affect the transmission of simple signals along simple motifs of molecular interaction networks. Even for very simple systems the effects of the different sources of variability alone and in combination can give rise to bewildering complexity. In particular extrinsic variability is apt to generate “apparent” information that can in extreme cases mask the actual information that for a single system would flow between the different molecular components making up cellular signalling pathways. We show how this artificial inflation in apparent information arises and how the effects of different types of noise alone and in combination can be understood.

## 1. Introduction

Information theory — as conceived by Claude Shannon — is the branch of the mathematical sciences that deals with the quantification of structures, regularities or semantic patterns in a stream of symbols or observations (Shannon, 1948). Information, different from meaning, was defined probabilistically in Shannon’s work and this notion has been applied with great success in the engineering and physical sciences (Peng *et al.*, 2005; Cover & Thomas, 2012). More recently, information theoretic approaches have gained in popularity in the biological sciences (Cheong *et al.*, 2011; Uda *et al.*, 2013).

It is obviously important for biological organisms, ranging from single cells to multi-cellular organisms, to sense, process and correctly adapt to their environment or their own physiological state (Endres & Wingreen, 2009; Andrews & Iglesias, 2007; Tkačik & Walczak, 2011; Tostevin *et al.*, 2012; Rhee *et al.*, 2012; Mc Mahon *et al.*, 2014). A host of recent studies have applied such information theoretical measures and analyses to biological systems, in particular gene regulation and signal transduction systems (Tkacik *et al.*, 2008b; Tostevin & ten Wolde, 2009; Tkacik *et al.*, 2012; Porter *et al.*, 2012; Selimkhanov *et al.*, 2014). In such a framework, we can for example, study how well a given molecular pathway relays information arriving at cell-receptors into the cellular interior, and in eukaryotes perhaps into the nucleus. If a cell “misinterprets” its environment — or fails to initiate an appropriate physiological response — then this can have obvious detrimental effects; we would therefore expect molecular signal transduction and information processing to have been finely honed by evolution. Information theory provides a framework in which we can attempt to quantify the accuracy and efficiency with which information is mapped onto physiological responses or actions (Tkacik *et al.*, 2008a; Doyle & Csete, 2011).

In molecular and cellular systems biology, but also in engineering or signal processing applications we are often primarily concerned with the efficiency of information transmission, which is also, of course, the context in which Shannon’s original work was set. For example, we may model the input and output of an information channel as random variables, *X* and *Z*, respectively. The *mutual information*, *I*(*X, Z*) is then a measure that tells us how much the uncertainty about *Z* is reduced if we know the state of the random variables *X* (Cover & Thomas, 2012). In terms of the entropies, *H*(*X*), *H*(*Z*) of random variables, *X, Z*, the canonical measure of the uncertainty associated to a random variable, we have

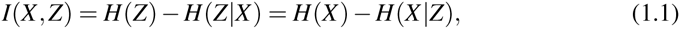

where *H*(*Z*|*X*) is the appropriate conditional entropy of *Z* if the state of *X* is known, *etc.*; below we assume that we can calculate entropies and derived quantities using the Lebesgue, counting or mixed measures as appropriate. For a perfect, noiseless channel information transmission is loss-less and, if the event spaces of *X* and *Z* are identical, we will therefore have, *H*(*Z*|*X*) = 0.

There are two subtle but we feel important differences between traditional applications of information theory and those found in biological systems. First, any molecular signal transduction pathway typically maps the input (such as an environmental stimulus), *X* onto an appropriate cellular response, (e.g. the concentration of an active transcription factor), Instead of faithfully reproducing the signal it is in fact *processed*, i.e. altered. Adaptive behaviour, where the response of the system attenuates back to the baseline level for a continuing stimulus, serves as a useful example, where *I*(*X, Z*) would vary over time and eventually, for perfectly adapted behaviour will approach zero. This can be partly accommodated into a conventional information theoretical approach by considering *X* and *Z* to be random variables with different event spaces (Tostevin *et al.*, 2012). This may happen, for example, for switch-like behaviour, where *Z ≈* 0 for *X* smaller than some threshold *X < X*_*t*_, and *Z ≈ c >* 0 for *X > X*_*t*_; here a continuous input is mapped onto distinct “ON” and “OFF” states (Tyson *et al.*, 2003).

The second difference lies in the physical manifestation of biomolecular signal transduction systems, which differ profoundly from their engineering counterparts: in physical systems there is a clear distinction between the channel or information transmission infrastructure (typically wires or antennas), the message (electrons or electromagnetic waves) and the energy required to deliver the message are typically distinct — although for e.g. single molecule transistors such a separation no longer holds. In biomolecular systems, the information processing machinery, the message, and the energy units are all molecules and medium and message are intricately linked (Kelly & Stumpf, 2008; Bowsher & Swain, 2014; Mc Mahon *et al.*, 2014).

Here we investigate how information is transduced by simple molecular systems, and using extensive simulation studies we attempt to distill the principles underlying molecular information processing. In particular we shall focus on noise and its impact on information processing in signal transduction (Bowsher & Swain, 2012; Komorowski *et al.*, 2013; Bowsher *et al.*, 2013; Selimkhanov *et al.*, 2014). The concept of “noisy channels” has been central to information theory since its conception (Shannon, 1948), but in a molecular context, as outlined above, we are not necessarily able to separate between the channel and the transmitted message.

Given that information is measured by entropy and that dynamics at the molecular scale tend to be stochastic we may expect systemic distortion of signals due to the noise inherent to molecular dynamics (Bowsher & Swain, 2012). Furthermore, in addition to such “intrinsic noise”, different individual cells will be subject to “extrinsic noise” sources (Swain *et al.*, 2002; Toni & Tidor, 2013). These include, by definition, variability among cells due to factors not explicitly considered in the analysis. In signal transduction this may, for example, be due to variability in the number of cell receptors, ribosomes, proteasomes, kinases, phosphatases *etc.*.

Below we will discuss the roles of intrinsic and extrinsic noise in the context of very simple signal transduction systems. We will focus on proteomic processes and very simple ON/OFF signals. Three main lessons emerge from this analysis: (i) noise, especially extrinsic noise, can lead to a systematic inflation in the apparent information transmitted through molecular interaction networks; (ii) transmission of stationary and even the simplest time-varying variable signals can differ quite profoundly (also in respect to which different noise sources affect the information transmission) even for very simple signalling motifs; and (iii) whereas we by now have good insights, often even intuition, as to how different molecular architectures/motifs affect the dynamics of a biological system, understanding the transmission of information, despite considerable recent progress, is still challenging. Even for the simple motifs considered here, a rich behaviour of even the conventional mutual information can be found.

## 2. Methods

In order to analyse the effects of noise in cellular signal processes we use information theoretical concepts, in particular mutual information. Obtaining its value is widely regarded as challenging and there is little consensus as to the best estimator of mutual information (Cellucci *et al.*, 2005; Fernandes & Gloor, 2010; Ziv *et al.*, 2007; Zhang & Zheng, 2015). Except for the special case where the joint distribution *p*_*X*,*Y*_ is multi-variate Gaussian, analytical values cannot be obtained. For this reason many different types of estimators have been devised; here we focus on a kernel density estimator based approach applied on instantaneous measurements of input and output states; we also use the linear noise approximation for trajectories, where we have analytical results as the distributions are multi-variate Normal (MVN).

### (a) Kernel Density Estimator

The kernel density estimation (KDE) was employed by Steuer *et al.* (2002). Considering a Gaussian kernel, a one-dimensional distribution *f* (*x*) is approximated from a data set 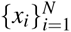 as

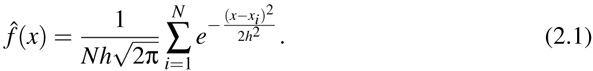

where *h* is the bandwidth. For our Gaussian kernel we use the Silvermann’s rule of thumb (Silverman, 1986), which is considered to be the optimal choice when the underlying distribution is Gaussian, 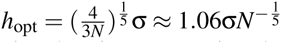, where σ is the standard deviation. These one and two dimensional estimates can be plugged into the continuous form of mutual information, which is a functional of probability densities

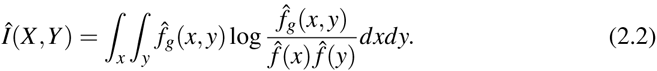

However, in order to simplify the algorithm we will typically represent the mutual information by

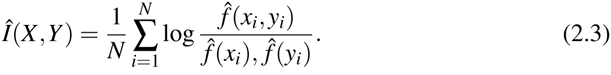

and sample *N* times from a mixture of multivariate Gaussians with mean (*x*_*i*_, *y*_*i*_) and apply the KDE with respect to each Gaussian. In addition we apply the copula transformation, by transforming the input data into quantiles (Nelsen, 2007). It is important to note that mutual information appears less sensitive with respect to the chosen smoothening parameter than the probability density.

### (b) The Linear Noise Approximation

When dealing with intracellular processes low copy numbers of molecules often lead to significant stochastic fluctuations governing the system dynamics. The Linear Noise Approximation (LNA) provides a reliable solution for many such systems, especially those for which the molecule number does not pass below *≈*10 and which are not highly non-linear (Wallace *et al.*, 2012). It has been previously successfully applied to simulate biochemical systems, for the inference of kinetic rate parameters (Komorowski *et al.*, 2009; Fange *et al.*, 2010), and for the sensitivity and robustness analysis of stochastic reaction systems(Komorowski *et al.*, 2011).

We consider a general system of *N* species made up of *X*_*i*_, *i* = 1,…, *N* molecules inside a volume Ω, giving a concentration *x*_*i*_ = *X*_*i*_/Ω. The state of the system can change by one of *R* chemical reactions corresponding to an event *j*, leading to a change in species *i* according to the stoichiometric matrix *S* = {*S*_*i j*_}_*i*=1,2,..*N*;_ *j*=1,2,*…R*. The probability of an event occurring in the time interval [*t, t* + *dt*] is given by the mesoscopic transition rates *ã*_*j*_(*x,* Ω, *t*). The LNA approximates the chemical master equation by dividing the system’s state into a deterministic and a stochastic part, which describe the the mean concentration of the reactants and the deviation of the reactant from their mean concentration values, respectively,

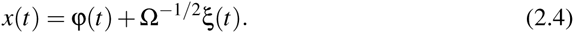

The equation describing the stochastic part is itself made up of two terms, respectively, comprising the drift *A* and diffusion *E*

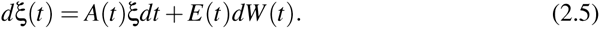

and the final distribution across all species is MVN at all times. We use the 

~~~
Stochsens
~~~

 package (Komorowski *et al.*, 2012) to obtain the LNA equations for our given systems.

### (c) Simulations

In addition to analyses using the LNA we also considered stochastic differential equations, which were simulated using the Euler-Maruyama approximation (Kloeden & Platen, 1992); the scipy library was used to solve ordinary differential equation. The simulations account for three different cases: the presence of intrinsic noise, of extrinsic noise, and of both. In order to account for extrinsic noise in the systems we consider parameters characterising different cells to distributed according to a Gaussian distribution. In our simulations we balanced the relative effects of extrinsic and intrinsic noise to study their respective effects on information transmission and simulations were used to calibrate the system such that the variance in system output at steady state due to each individual noise source amounts to 10% of the mean/deterministic steady state abundance.

## 3. Results

### (a) Noise in Simple Motifs

We begin by estimating the mutual information between molecular species that form the inputs and outputs of three increasingly complex molecular signalling motifs(Alon, 2007; Ingram *et al.*, 2006; Domedel-Puig *et al.*, 2010) under the effects of different types of noise, see Fig. 1. The first motif we consider is a basic input-output system with just two molecular species, corresponding to, for instance, a kinase and its regulated target substrate, with no additional interactions. Next, we introduce an additional species, to create a cascade motif, made up of a chain of elements with linear dependencies, which in a biological context could represent a simple signalling cascade (Miller *et al.*, 2008). The final motif is a three-species system of the simple feed-forward loop (FFL) type (Mangan & Alon, 2003; Alon, 2007). Note that there are eight different structural types of FFL based on different combinations of activation and repression, each categorised into coherent and incoherent depending on whether the sign of the direct and indirect regulation path are the same or opposite. Here the focus is on the so-called coherent type-1 FFL which appears commonly in both *E. coli* and *S. cerevisiae* (Mangan & Alon, 2003).

**Figure 1.**
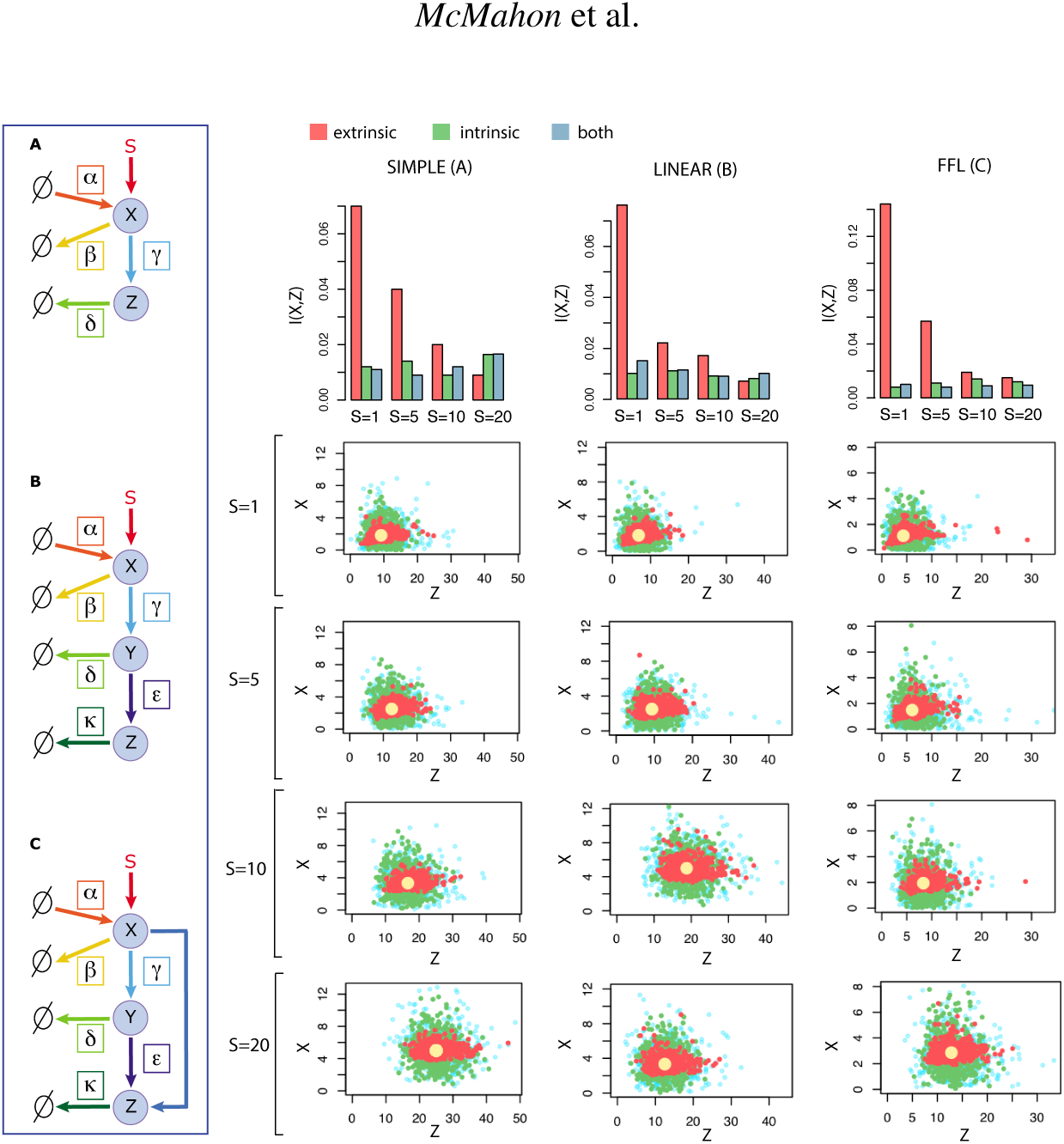
Here we show the behaviour of the simple input-output motif (A), the linear motif (B) and the feed forward loop motif (C), in the presence of extrinsic (orange), intrinsic(green), and both types of noise (blue) — all with comparable output variances — as well as in the absence of noise (yellow). For simplicity all three motifs contain a single time-varying stimulus *S*(*t*) and molecules with fixed degradation rates. The scatter plots depict the distributions of species *X* and *Z* for each motif (simple, linear and feed forward loop from left to right), and show how the molecular species react to different input signals *S* = {1, 5, 10, 20}, increasing from the top to the bottom rom respectively. The bar charts represent the trend in mutual information computed via KDE between molecular species *X* and *Z* for the motifs in the presence of the above mentioned different input signals.

Using a stochastic differential equation (SDE) model, extrinsic noise is introduced via parameters that follow themselves a Gaussian distribution. We observe that most of the time the mutual information *I*(*X*; *Z*) is highest for the cases with extrinsic-only noise; information transmission is most affected, therefore, by the presence of intrinsic noise and, perhaps counter-intuitively, the loss in fidelity is not sufficiently mitigated by increasing signal strengths. Interestingly the mutual information value in the presence of both types of noise appears, most of the time, to be between that displayed solely in the presence of intrinsic and solely extrinsic types of noise. This is somewhat counter-intuitive as we would expect the mutual information of the systems in the presence of both types of noise to be the lowest of the three. It leads us to believe that the two types of noise interact in a non-additive manner.

Furthermore, although this conclusion about the relative impact of the different types of noise cannot be generalised to all signalling pathways, the observed pattern is surprisingly similar across the three otherwise non-trivially different systems. The pattern also remains consistent when analysing these systems with different input signals *S*. In addition, our results show that increasing the value of *S* decreases the mutual information significantly in the presence of extrinsic noise.

To investigate the generality of our results further, the same set of motifs was also considered but accounting for the possibility of basal transcription and activation of each of the molecular species under mass action kinetics — this would be expected to reduce the ability of the system to trace the signal appropriately. In Figure 2 we show the trajectories for molecular species *Z* in the simple, linear and feed forward loop obtained with the LNA compared to 100 trajectories obtained with the SDE; the mean value of the latter (red line) corresponds to the ODE trajectory obtained with the former (green line). As is apparent in the figure, while the average behaviour of the LNA and the SDE are in excellent agreement (as are their respective variances), the (analytical) MI estimates obtained for the LNA are consistently higher than those obtained from the SDE simulations (which were estimated using KDE described in the *Methods* section); this reflects the way in which the LNA fails to capture non-Gaussian noise. The MI estimates in figure 2 for the LNA appear inflated compared to the SDE case, as the LNA restricts the joint distribution of the state variables, here input, *X*, and output, *Z*, compared to the SDE (which captures the stochasticity of the system fully). Nevertheless both estimates display the same qualitative dependence on the signal and are consistent across motifs.

**Figure 2.**
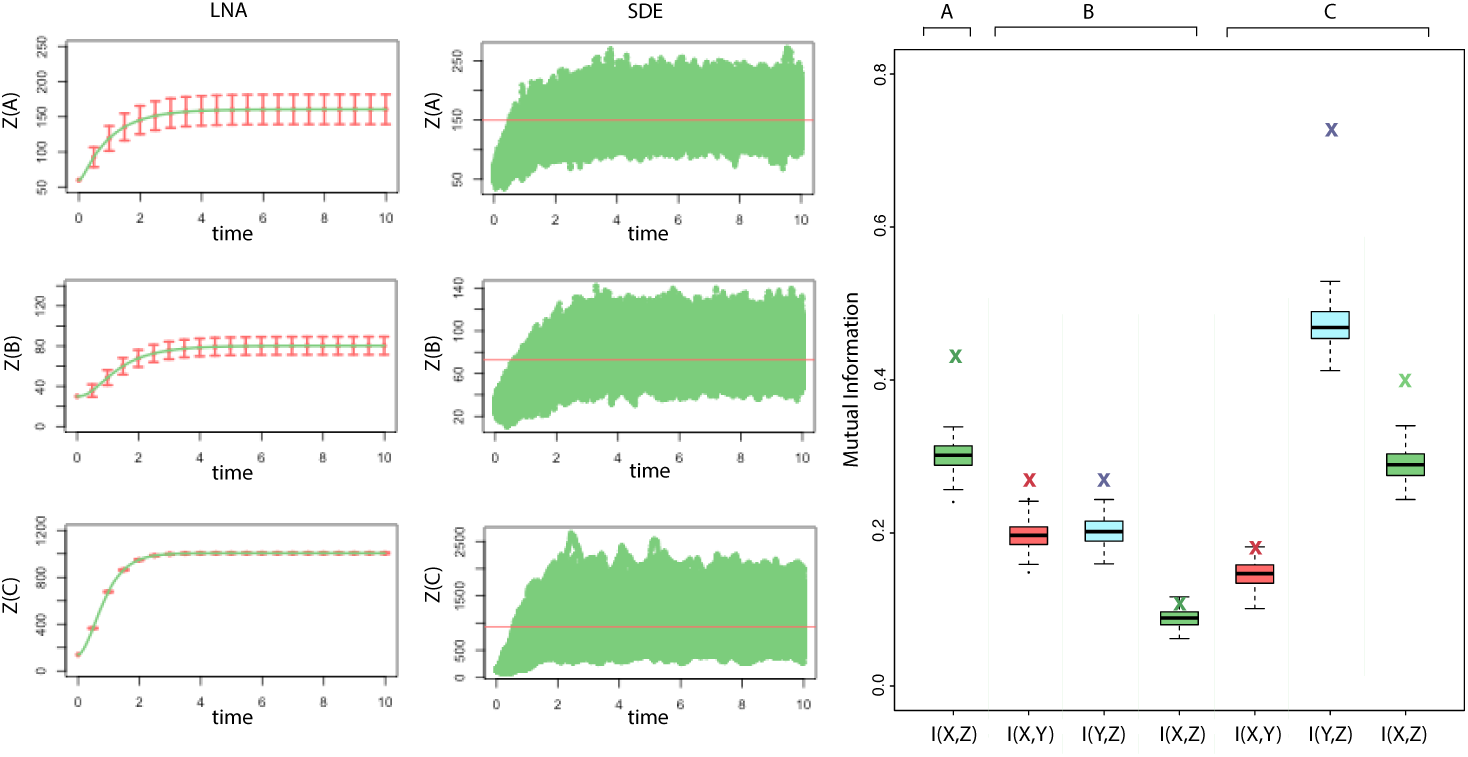
Here we show the trajectories for molecular species *Z* simulated with both LNA and SDE, for the simple (A), linear (B), and feed forward loop (C) motifs. The box plot displays the mutual information for these motifs computed analytically with the LNA based approach (displayed by a cross symbol) compared to 100 estimates obtained with the kernel density estimator which employs the SDE results. While the LNA is capable of capturing the average behaviour, the variability of the output is reduced considerably compared to the SDE case, and this is reflected in the apparent increase in mutual information observed for the LNA compared to the SDE case.

Thus far we have only looked at stationary signals. Biologically more interesting are, of course, dynamically changing signals — such as spatial differences in nutrient abundance or temporally varying environmental signals. For simplicity we consider a very simple form for a signal that changes with time: a square wave process that alternates between successive “ON” and “OFF” states. We model the motifs under two scenarios of this signal, corresponding to different switching rates, and investigate the covariances and mutual information values between the inputs and outputs of the three motifs for different levels of noise.

In a first instance, the motifs are simulated with the LNA which already accounts for the intrinsic noise, to which varying degrees of extrinsic noise are added as displayed in Figure 3. it was interesting to see how the mutual information varies in relation to the signal dynamics. In fact it oscillates between two values, reaching its peak and base points just after the switch is turned OFF and ON respectively. The average mutual information between input and output is not affected by the level of extrinsic noise, but its variance increases quite considerably; we do, however, observe that this increase with extrinsic noise is somewhat suppressed as the motifs become more complicated (as the effects of different origins of noise can balance one another out). Interestingly, the covariances shown in the figure trace the dynamics of the inputs more faithfully than do the MI estimates; this is because the MI also depends on the individual variances of *X* and *Z*, which have their own temporal dependencies. The details of this do, of course, depend on the parameters of the motifs analysed here, but the results obtained here are characteristics for the behaviour that can be observed for even such simple motifs. Obviously, signalling dynamics will depend on the frequency characteristics of the input signal as well.

**Figure 3.**
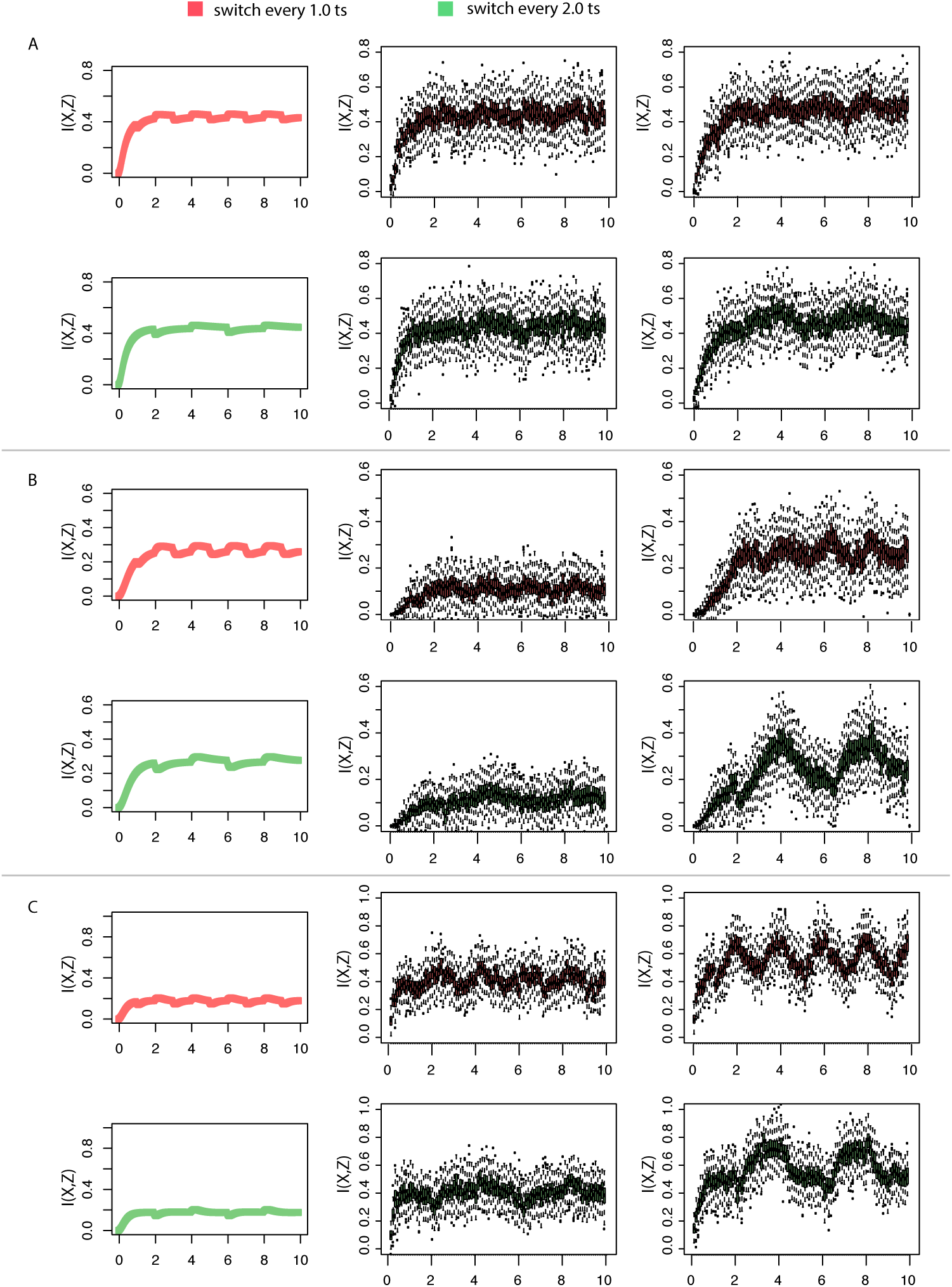
The figure displays the mutual information values across time between variables *X* and *Z*,in the simple, linear and feed forward motif (from top to bottom respectively) in the presence of different noise perturbations. The first column represents the system in the presence of intrinsic noise only, already included in the linear noise approximation simulation; as we go from left to right we gradually increase the amount of added extrinsic perturbation. The simulations were performed in the presence of a signal *S* switching between an on and off state *S* = [0, 1], every 1.0 (red) or 2.0 (green) time steps (the integration step used was *dt* = 0.01).

In particular, for the FFL, depending on the parameters, any number of different types of behaviour can be observed. But generally, we find that the mutual information between *X* and *Z*, *I*(*X, Z*), is less variable, compared to the other motifs, as the extrinsic noise is increased. More generally, the mutual information traces the signal more faithfully for the FFL, as *X* affects *Z* both directly and indirectly via *Y*, which integrates out some of the variability resulting from extrinsic noise.

To complete the analysis of the motifs under dynamic conditions, we analyse the behaviour of the motifs with the same signal conditions, simulated with ordinary differential equations but perturbed by extrinsic noise only, and compared them to the corresponding stochastic system (intrinsic noise) and a system with both types of noise. In figure 4 we show trajectories for molecular species *Z* in the three systems. What can be seen from the trajectories, is that the intrinsic noise appears to govern the dynamics of *Z*; in fact, combining both noise sources affects the trajectories only marginally. In the lower part of the figure we focus on the mutual information estimates. The mutual information was computed via the KDE across all types of noise for two specific time points. The time points, *T* 1 and *T* 2 displayed on the trajectories of species *Z*, were selected based on the observations made in 3; specifically we chose time points just just after the switch is turned OFF and ON to estimate the peak and base in the mutual information oscillation.

**Figure 4.**
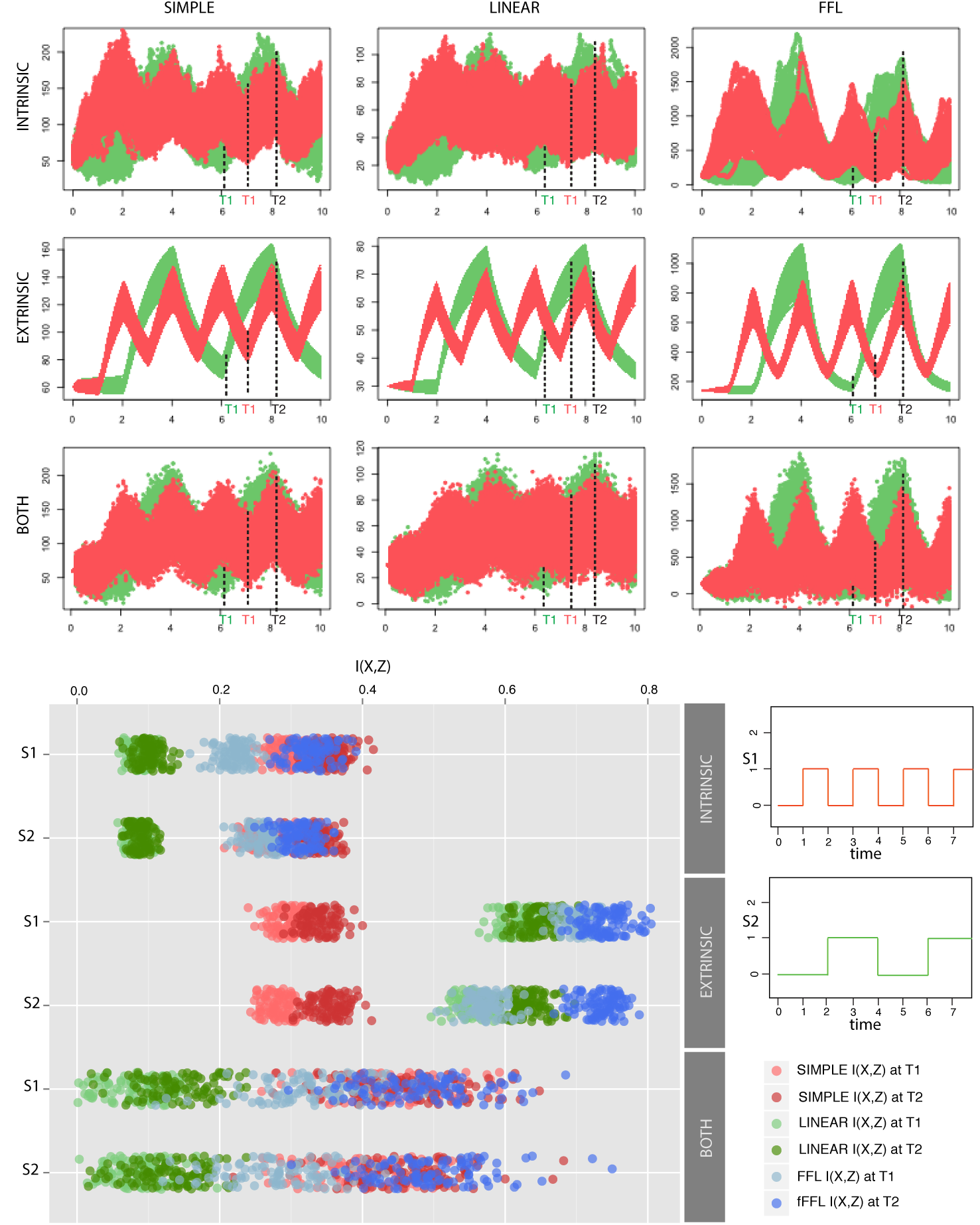
The figure displays the trajectories of species *Z*,in the simple input-output, linear and feed forward motif in the presence of different noise perturbations, intrinsic, both, and extrinsic, from top to bottom respectively. The systems were simulated for different signal frequencies *S*1 and *S*2, displayed in the bottom right side of the figure. For each of the frequencies different time points T1 and a time point T2 were selected at which to estimate the mutual information. The strip chart displays the mutual information values computed via the KDE between species *X* and *Z* for the systems in question, under each type of noise conditions.

We show the estimated mutual information for the linear three-node motif and the FFL are highest for the ODE with extrinsic noise, while for the simple motif it is lowest. This result is highly dependent on the relative sizes of the different parameters and can be explained by the effect that different parameters have on the system dynamics. Generally, cells that have different internal parameters will map inputs onto different outputs, which may appear as inflated information transmission: note that the steady-state abundances (if steady states exist) of all molecules will depend on the parameters. We also compare the information transmission efficiency for different signal frequencies (*S*1 and *S*2); the effects here are more pronounced in the presence of both sources of noise: when extrinsic noise is present we find a greater difference between minimal and maximal transmitted mutual information. When only intrinsic noise is present, this apparent dependence of transmitted information on the frequency of the signal is not pronounced. Across these systems it would appear once again that the addition of intrinsic noise to extrinsic noise is not cumulative, as the value of mutual information in the presence of both types of noise seems to be oscillating around that of the intrinsic noise.

### (b) Noise in protein expression and activation

So far we have considered generic models that have previously been described in, or applied to, biological signalling or regulation dynamics. Here we apply the same perspective to a model of protein expression that is more immediately connected to biological processes (Elowitz *et al.*, 2002; Ingram *et al.*, 2008). Protein expression requires a cascade of biomolecular reactions to produce functional protein. Each reaction is associated with a relative loss of information, but some may distort signals more than others. We consider the model used by Komorowski *et al.* (2013) to describe gene expression and activation of the protein product via reversible phosphorylation, where the kinase and phosphatase are assumed to be abundant and at constant activity levels.

The following equations, involving mRNA *m*, protein, *P* and active (phosphorylated) protein, *P\**, were considered in order to simulate the system in a similar fashion as above:

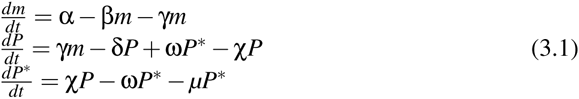

The model represented in Figure 5, was considered with different rates of dephosphorylation (parameter ω) and degradation of the active protein (parameter *μ*), which were previously shown to be the reactions that make the largest relative contributions to the variability in the abundance of the active protein(Komorowski *et al.*, 2013). For this system we proceed as before and estimate the MI for the three noise scenarios between the three molecular species at steady state. Again we find that extrinsic noise leads to an apparent increase in the mutual information, see Figure 5, whereas intrinsic noise leads to a clear reduction in the mutual information. Mutual information is always highest between *P* and *P\** but low overall. Again we observe a trend where mutual information appears to increase in the presence of extrinsic noise and in the presence of both types of noise the mutual information typically takes on intermediate values.

**Figure 5.**
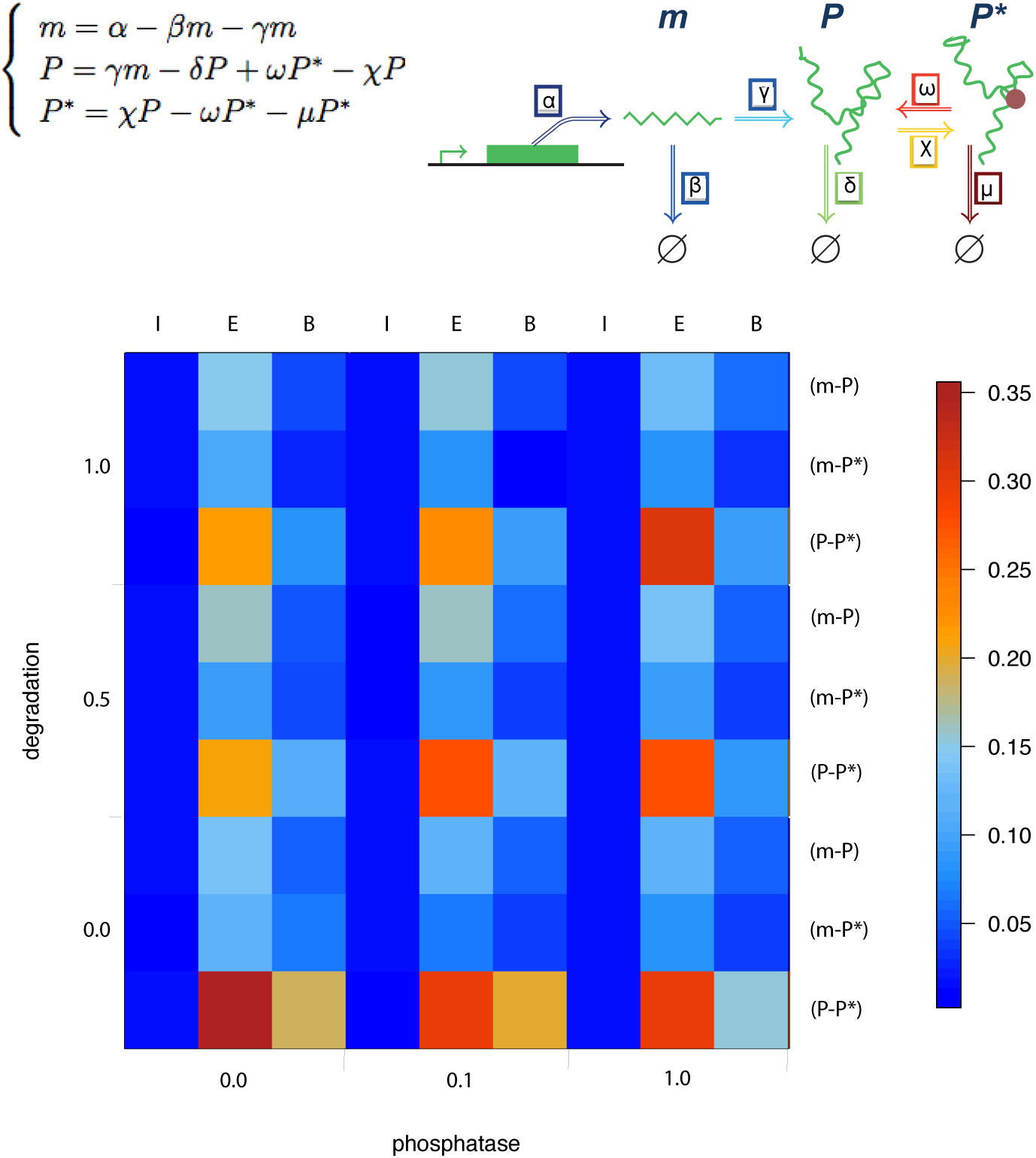
Heat map representation of the mutual information values in relation to different phosphatase and degradation activity. We consider different rates of dephosphorylation (phosphatase activity) and protein degradation and find for extrinsic noise that when these rates are minimal, the mutual information between mRNA, *m*, protein *P*, and active protein *P\** becomes maximal. It decreases as the rates are increased, but increases again as both the degradation and phosphatase activity increase. Intrinsic noise alone results in negligible transmission of information irrespective of the rate constants for degradation and dephosphorylation.

The dependence of the mutual information (for the two cases exhibiting extrinsic noise) on the rates of dephosphorylation and degradation is such that it initially decreases as the rates of these two processes increase, before increasing again with further increase in the dephosphorylation and degradation rates. We will revisit these observations below. The most important result of this finding is that the apparent amount of mutual information between e.g. mRNA and protein or active protein can be inflated by cell-to-cell variability due to extrinsic sources of variability.

## 4. Discussion

An important issue to remember is that while mutual information can shed light upon the effectiveness of transmission of information, information is only a statistical measure for the regularity of patterns in a stream of data — not all of this may be biologically relevant. Extrinsic noise — the systematic differences in molecular parameters between different cells — will often (but not always) act to distort or stretch out signals (see also figure 6). For the purpose of illustration we consider a system with two molecular species described by random variables *X* and *Y* (e.g. the simple linear motif), with *x* ∼ *f* (*S,* θ) and *Y ∼ g*(*x,* θ) (we consider systems where *Y* is independent of *S* conditional on the state of *X*). Then given an input signal (which may change over time *S*(*t*), in cells which have identical parameters θ_0_ we obtain measurements *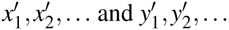,*

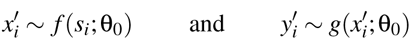

In the presence of extrinsic noise the cells will differ in their respective parameters and we obtain *x*_1_, *x*_2_, *…* and *y*_1_, *y*_2_, *…*,

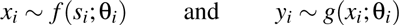

Then is we generally expect (and have indeed found in the results shown here) that

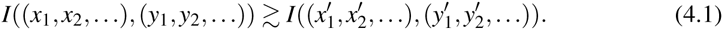

The dynamics of the signal transduction system affect the mutual information as well as the entropies of the random variables *X* and *Y*; suppression of the effects of extrinsic noise will not be the rule, but its effect will be reduced is intrinsic noise is appreciable. We can also rationalize the inequality (4.1) by considering the effects of extrinsic noise on the terms in the definition of the mutual information:

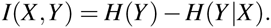

Extrinsic noise will tend to lead to an increase in the spread of *Y* and hence *H*(*Y*) will increase under extrinsic noise. Depending on the dynamics of the signalling system we would also expect the conditional entropy *H*(*Y|X*) to decrease as both *Y* and *X* are functions of the parameters θ that differ between cells.

For intrinsic noise alone the interplay between the dynamics of the molecular information processing system and the concomitant inherent stochasticity are already difficult enough to disentangle and have attracted considerable attention (Elowitz *et al.*, 2002; Cai *et al.*, 2006; Friedman *et al.*, 2006; Ingram *et al.*, 2008). Especially for non-linear systems the combined effects of noise and dynamics can give rise to rich and diverse behaviour of the system (Bowsher *et al.*, 2013). Here we have mostly focussed on the stationary dynamics and there, as far as the information transmission is concerned, we can typically ignore much of this complexity (provided a stable set of equilibrium solutions exists). Because of the lack of normalization of the mutual information, the channel capacity (Cover & Thomas, 2012; Uda *et al.*, 2013),

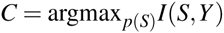

is sometimes preferred over the mutual information; this is a variational problem over the possible input distributions, *p*(*S*), of the signal, *S*.

But the channel capacity also implicitly depends on the parameters, θ, characterizing the information processing network, *i.e.* the function, *f* (*S*; θ). This makes the interpretation of the information processing capability of populations of systems/cells in the presence of extrinsic noise less straightforward. In principle we could consider the channel capacity averaged across the ensemble but this would hopelessly skew the results, as it will be the between-cell variability that will drive the “apparent” information between inputs and outputs that is captured by the mutual information. In each single cell — or any ensemble of cells with the same kinetic parameters — the mutual information will be much smaller as intrinsic noise alone will only decrease and never increase information — of course, extrinsic noise only increases apparent information (the “level of surprise” at seeing a given symbol/signal). Taken together we cannot predict *a priori* how these contrasting forces will interact. Certainly the effects of different types of noise on signal transduction are not simply additive.

Extrinsic noise, *i.e.* different parameters characterising the biomolecular reaction networks in different cells, can even lead to qualitatively different behaviour across a population of cells; some cells might, for example, oscillate, while others attain a stable equilibrium (depending on the eigenvalues of the corresponding Jacobian matrices describing the different systems). Different parameters, see figure 6, are associated with different gradient fields that may drive solutions (even for identical initial values) to diverge; especially for linear and monotonic systems we would then expect to observe an apparent increase in the mutual information for extrinsic uncertainty. Intrinsic noise, by contrast will typically (but not always) broaden a deterministic solution. How and when these different sources of noise work together and how they affect information transmission, is highly dependent on the system under consideration; this is especially true for non-linear and non-monotonic dynamical systems.

However, it is clear that these types of noise do not contribute to the information transmission across the system in a simple additive way. The present analysis appears to suggest that extrinsic noise can give rise to “apparent information” and it is important to be aware of this when assessing biological information processing systems, or comparing single-cell and population level processes from an information theoretical perspective.

**Figure 6.**
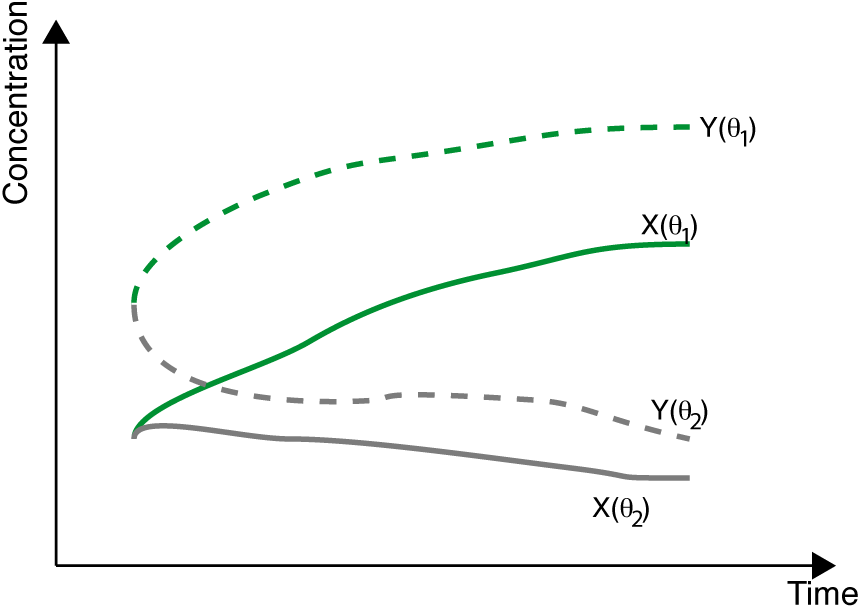
For suitable dynamical systems, differences in parameters (due to extrinsic noise) will give rise to diverging solutions, (*X* (θ),*Y* (θ)) and the differences between θ and θ *′* may suffice to drive differences in estimates of the mutual information between *X* and *Y*.

What this analysis has provided is a quantitative assessment of the effects of different types of noise on the information transmission along simple network motifs inspired by biophysical systems. We feel that there are two important lessons that follow from this work: (i) even for very simple systems and simple signals the information theoretic analysis reveals rich and diverse behaviour. Because of the statistical definitions of entropies and mutual information this diversity may be hard to glean from looking at the dynamics of the system alone; instead we really have to understand the effect of the dynamics of the molecular reaction network onto the distributions of inputs; (ii) as single cell data are becoming available more routinely it becomes important to be able to deal with extrinsic noise as it tends to affect our assessment of biological information processing and can lead to inflated estimates of the mutual information from single cell data. There are different ways to implement extrinsic noise and the one chosen here is perhaps among the most straightforward and convenient (Swain *et al.*, 2002; Toni & Tidor, 2013).

The comparison of the LNA with exact stochastic simulations was instructive in showing that the amount of variability and the shape of the distribution of outputs can play a profound role. The LNA may miss some of this as real distributions may differ quite substantially from the underlying Gaussian assumption in the LNA, especially for low molecule abundances and/or non-linear dynamics. The dynamical features of a system affect the information transmission even at stationarity. Analysis of such simple motifs and the way that they shape cellular information transmission can only be a first step, but it is a necessary one, towards understanding of cellular decision making processes. Relevant information is, however, not always just encoded in terms of abundances (or amplitudes); frequency and gradients are sensed as well and for these more explicitly dynamical notions of mutual information such as the transfer entropy need to be considered from the outset. The present results suggest, however, that the role of extrinsic noise ought to be considered explicitly, as failure to properly account for its effects is likely to be misleading.

SMM and OL acknowledge studentship support from the Department of Life Sciences and the BBSRC, respectively. MPHS is a Royal Society Wolfson Research Merit Award holder and acknowledges support from the BBSRC, EPSRC and the Leverhulme Trust. We thank the members of the *Theoretical Systems Biology Group* for helpful discussions.

